# MNK inhibitor eFT508 (Tomivosertib) suppresses ectopic activity in human dorsal root ganglion neurons from dermatomes with radicular neuropathic pain

**DOI:** 10.1101/2023.06.13.544811

**Authors:** Yan Li, Megan L. Uhelski, Robert Y. North, Juliet M. Mwirigi, Claudio E. Tatsui, Juan P. Cata, German Corrales, Theodore J. Price, Patrick M. Dougherty

## Abstract

Spontaneous activity in dorsal root ganglion (DRG) neurons is a key driver of neuropathic pain in preclinical models and in patients suffering from this largely untreated disease. While many intracellular signaling mechanisms have been examined in preclinical models that drive this spontaneous activity (SA), none of these have been tested directly on spontaneously active human nociceptors. Using cultured DRG neurons recovered during thoracic vertebrectomy surgeries, we show that inhibition of mitogen activated protein kinase interacting kinase (MNK) with eFT508 (25 nM) reverses SA in human sensory neurons associated with painful dermatomes. MNK inhibition in spontaneously active nociceptors decreased action potential amplitude and produced alterations in the magnitude of afterhyperpolarizing currents suggesting modification of Na^+^ and K^+^ channel activity downstream of MNK inhibition. The effects of MNK inhibition on SA took minutes to emerge and were reversible over time with eFT508 washout. MNK inhibition with eFT508 led to a profound loss of eIF4E Serine 209 phosphorylation, a specific target of the kinase, within 2 min of drug treatment, consistent with the rapid action of the drug on SA in electrophysiology experiments. Our results create a compelling case for the future testing of MNK inhibitors in clinical trials for neuropathic pain.

**Conflict of interest:** TJP is a co-founder of 4E Therapeutics, a company developing MNK inhibitors for neuropathic pain. The other authors declare no conflicts of interest.

## Introduction

Neuropathic pain is largely untreated with existing therapeutics and is a leading cause of disability and suffering world-wide. Neuropathic pain can be alleviated with local or neuraxial nerve blocks in most patients, demonstrating a peripheral driver for pain (Haroutounian et al., 2014). Preclinical neuropathic pain models are associated with emergence of SA in sensory neurons, including nociceptors (Walters et al., 2023). Microneurography studies have demonstrated the presence of SA in peripheral nociceptor axons in neuropathic pain patients (Serra et al., 2012), and our recent work has demonstrated a direct association between SA recorded *in vitro* and the presence of neuropathic pain in associated dermatomes in thoracic vertebrectomy surgery patients with neuropathic pain (North et al., 2019). Collectively, these lines of evidence strongly support nociceptor SA as a key driver of neuropathic pain, in particular the spontaneous pains that are characteristic of so many neuropathic pain disorders.

A candidate intracellular signaling mechanism underlying generation of SA in nociceptors is MNK signaling. MNK1/2 are downstream targets of ERK and p38 and both genetic and pharmacological inhibition of MNK signaling reduces evoked and spontaneous aspects of neuropathic and other forms of pain in animal models (Moy et al., 2017; Megat et al., 2019; Mihail et al., 2019; Barragan-Iglesias et al., 2020; Shiers et al., 2020; Lackovic et al., 2023). The MNK inhibitor eFT508 has recently been named Tomivosertib because this drug has entered late-stage clinical trials for oncology indications (Stumpf et al., 2018). The specificity of this molecule, as well as its demonstrated safety profile in humans makes it an ideal candidate for testing the hypothesis that MNK signaling plays a key role in generating SA in human nociceptors. Our findings support the development of MNK inhibitors for neuropathic pain.

## Methods

### Human DRG study approval

Human tissue procurement procedures were approved by the Institutional Review Board at the University of Texas MD Anderson Cancer Center and the University of Texas at Dallas and all experiments conformed to relevant guidelines and regulations. Written informed consent for participation, including use of tissue samples, was obtained prior to inclusion. Consent for research was obtained from next of kin by the Southwestern Transplant Alliance (STA) for organ donors. Tissue samples from a total of 11 patients were used in the study (see Supplemental Table 1).

### Human dorsal root ganglion neuron preparation

Human DRG neurons were prepared for electrophysiology as described previously (Li et al., 2015; Li et al., 2018). Briefly, DRG tissue was obtained from patients undergoing surgical treatment that necessitated ligation of spinal nerve roots to facilitate tumor resection and/or spinal reconstruction or were collected from organ donors. Excised DRG were transferred immediately into cold (∼4°C) and sterile balanced salt solution (EBSS, gibco, Cat# 24010-043) and transported to the laboratory on ice in sterile, sealed 50-ml centrifuge tubes. Each ganglion was then carefully dissected from the surrounding connective tissues and sectioned. Sections of DRG were divided into 1 mm^2^ sections using a surgical scalpel and placed in a Petri-dish containing trypsin (Sigma, Cat# T9201), collagenase (Sigma, Cat# C1764), and DNase (Sigma, Cat# D5025) in DMEM/F12 (gibco, Cat# 11330-032) and placed in an incubated orbital shaker at 37°C for 20 min. After each 20 min session, the digestion solution was collected, placed into a blocking solution (DMEM/F12 with 10% horse serum, gibco, Cat# 26050-070), and replaced with fresh digestion solution. This process was repeated until the sections were adequately digested. The collected digestion solution and blocking solution was centrifuged at 23°C and 180g for 5 min. The supernatant was removed, and the cell pellet re-suspended with 1 mL of culture media consisting of DMEM/F12 (gibco, Cat# 11320-033) with 10% horse serum, 1% penicillin-streptomycin (gibco, Cat# 15140-122) and 0.1 ug/mL nerve growth factor (NGF, Cell signaling Technologies, Cat# 84087S). An additional 2-3 mL of culture media was added, and then the suspension was filtered through a 100 µm cell strainer. The remaining suspension was centrifuged at 23°C and 180*xg* for 5 min. The supernatant was removed, and the cell pellet re-suspended with culture media. Cells were plated on glass sheets coated with poly-L-lysine (Sigma, P2636) and held in culture dishes with culture media until used (Li et al., 2018).

For immunostaining experiments, lumbar DRGs were harvested from a 50-year old female donor approximately 3 h after cross-clamp and placed in chilled, oxygenated artificial cerebrospinal fluid (aCSF) prepared using a protocol adapted from Valtcheva et al., 2016(Valtcheva et al., 2016). Sterilized 15mm glass coverslips (Chemglass Life Sciences, Cat# CLS1760015) were coated with 0.01 mg/mL poly-D-lysine (Sigma, Cat# P7405-5MG) on Falcon® 12-well plates (Sigma-Aldrich, Cat#SIAL0513-50EA) overnight at 4°C and stored up to two weeks in wrapped aluminum foil. The recovered DRG was trimmed with No. 5 forceps (Fine Science Tools, Cat#11252-00) and Bonn scissors (Fine Science Tools, Cat# 14184-09) to expose the cluster of cell bodies. At the time of culturing, a final enzyme solution consisting of 2mg/mL Stemzyme I (Worthington Biochemical, Cat#LS004106) and 0.1 mg/mL DNAse I was prepared in HBSS (Thermo Scientific, Cat#14170161) and allowed to incubate in a water bath. A total of 5 mL of enzyme solution was used to dissociate the DRG tissue. Ganglia were finely minced with scissors into smaller fragments less than 3 mm. The tissue fragments were transferred to the prewarmed enzyme solutions in 10 mL conical tubes and placed in a 37°C shaking water bath. The tissue was regularly triturated every 25 minutes using a fire-polished sterile glass Pasteur pipette (VWR, Cat#14673-043) until the tissue completely dissociated resulting in a cloudy solution (approximately 2 hrs in total). The solution containing dissociated neurons was passed through a 100 µm sterile cell strainer (VWR, Cat# 21008-950) into a 50 mL conical tube. The cell suspension was then gently added to a 3 mL 10% Bovine Serum Albumin (Biopharm, Cat#71-040)/HBSS gradient and spun down at 900*xg* for 5 min at RTP, 9 acceleration, 5 deceleration. The resulting supernatant was resuspended in prewarmed and sterile-filtered DRG media comprising of Brainphys® media (STEMCELL Technologies Cat#05790) supplemented with 1% Pen/Strep (Thermo Scientific, Cat#15070063), 1% GlutaMAX® (United States Biological, Cat#235242), 2% NeuroCult™ SM1 (STEMCELL Technologies, Cat# 05711), 1% N-2 Supplement (Thermo Scientific, Cat#17502048), 2% HyClone™ Fetal Bovine Serum (ThermoFisher Scientific Cat#SH3008803IR), 10 ng/mL human β Nerve Growth Factor (Cell Signaling Technology, Cat# 5221SC), and 3 μg/mL + 7 μg/mL FRD+U (5-Fluoro-2′-deoxyuridine, Sigma-Aldrich, Cat# F0503 and Uridine, Sigma-Aldrich, Cat# U3003-5G). 120 µL of cell suspension was carefully plated onto each coverslip and allowed to adhere for 2 hours at 37°C and 5% CO_2_. The wells were flooded with 1500 µL prewarmed complete DRG media. The cell cultures were maintained for 5 days prior to use. Media changes were performed every other day.

### Whole cell recording of spontaneous action potential in acutely dissociated rat DRG neurons

The day after plating, dissociated DRG neurons were transferred to a recording chamber perfused with oxygenated (95% O_2_ + 5% CO_2_) extracellular solution containing 117 mM NaCl, 3.6 mM KCl, 1.2 mM NaH_2_PO_4_ H_2_O, 2.5 mM CaCl_2_, 1.2 mM MgCl_2_, 25 mM NaHCO_3_ and 11 mM glucose adjusted to pH 7.4 with NaOH. Glass-micropipettes were filled with an internal solution with 125 mM KCl, 15 mM K-gluconate, 5 mM Mg-ATP, 0.5 mM Na_2_GTP, 5 mM HEPES, 2 mM MgCl_2_, 5 mM EGTA, and 0.5 mM CaCl_2_ adjusted to pH 7.4 with KOH. The DRG neurons were then held at 0 pA to record spontaneous action potential for 5 min. DRG neurons were then held at 0 pA, and action potentials were evoked using a series of 500-ms depolarizing current injections in 10-pA steps from -50 pA. The current that induced the first action potential was defined as the current threshold (rheobase). Only DRG neurons with a resting membrane potential of at least -40 mV, stable baseline recordings, and evoked spikes that overshot 0 mV were used for further experiments and analysis. Series resistance (Rs) was compensated to above 70%. All recordings were made at room temperature. eFT508 was purchased from MedChem Express (Cat# HY-100022).

### Immunocytochemistry and imaging

The treated cultures were immediately fixed with 10% formalin (ThermoFisher Scientific, Cat# 23-245684) for 15 min at RT and rinsed three times with 1X PBS. 1 h block was performed at RT with 10% NGS and 0.3% Triton X-100 in 1X PBS. The coverslips were then incubated overnight at 4°C with primary antibodies chicken anti-peripherin (1:1K, EnCor Biotechnology, Cat# cpca-peri) and rabbit anti-p-eIF4E (1:1000, phospho-S209, Abcam, Cat# ab76256). Coverslips were rinsed 3 times with 1X PBS for 15 min and incubated with secondary antibodies goat anti-chicken IgY (1:2K, H+L, Alexa Fluor 647, Invitrogen, Cat# A21449) and goat anti-rabbit IgG (1:2K, H+L, Alexa Fluor 555, Thermo Fisher Scientific, Cat# A21428) for 1 h at RT. The coverslips were washed 3 times with 1X PBS and mounted with prolong gold onto uncharged glass slides and allowed to cure overnight. Slides were imaged at 20X magnification using the Olympus FV3000 RS confocal laser scanning microscope.

### Data Analysis

For electrophysiology experiments, data were expressed as the mean ± standard error of mean. Results were analyzed using non-parametric Wilcoxon matched-pairs signed rank tests or two-way ANOVA. For immunostaining experiments, images were analyzed using Olympus cellSens software version 1.18. Mean gray intensity values for each region of interest were normalized to the control group at each timepoint and plotted. Two-way ANOVA with Bonferroni multiple comparisons was performed to assess differences between groups. P < 0.05 was considered statistically significant. Statistical tests were done using Prism 9.5.1 software program (GraphPad Software).

## Results

### eFT508 suppresses SA in human sensory neurons associated with painful dermatomes

We conducted whole cell patch clamp experiments on human DRG neurons cultured after recovery from thoracic vertebrectomy surgeries. Using DRGs associated with painful dermatomes we recorded from cells that exhibited clear SA and then treated cells with either vehicle (DMSO 0.0025%) or eFT508 (25 nM, in the same concentration of DMSO). While vehicle treatment had no effect on SA over the recording period (**Fig 1A**), eFT508 decreased spontaneous activity in a fashion that required 5-10 min of treatment prior to observation of a decrease of SA and that was reversed upon drug washout after a 5-10 min recovery period (**Fig 1B and C**). In additional recordings we tested the effect of eFT508 on step voltage-evoked current changes and observed no clear effect on any specific aspect of the whole cell current-voltage relationship in all neurons grouped together (**Fig 1D**) or specifically in neurons from male or female patients (**Fig 1E and F**).

**Figure 1.**
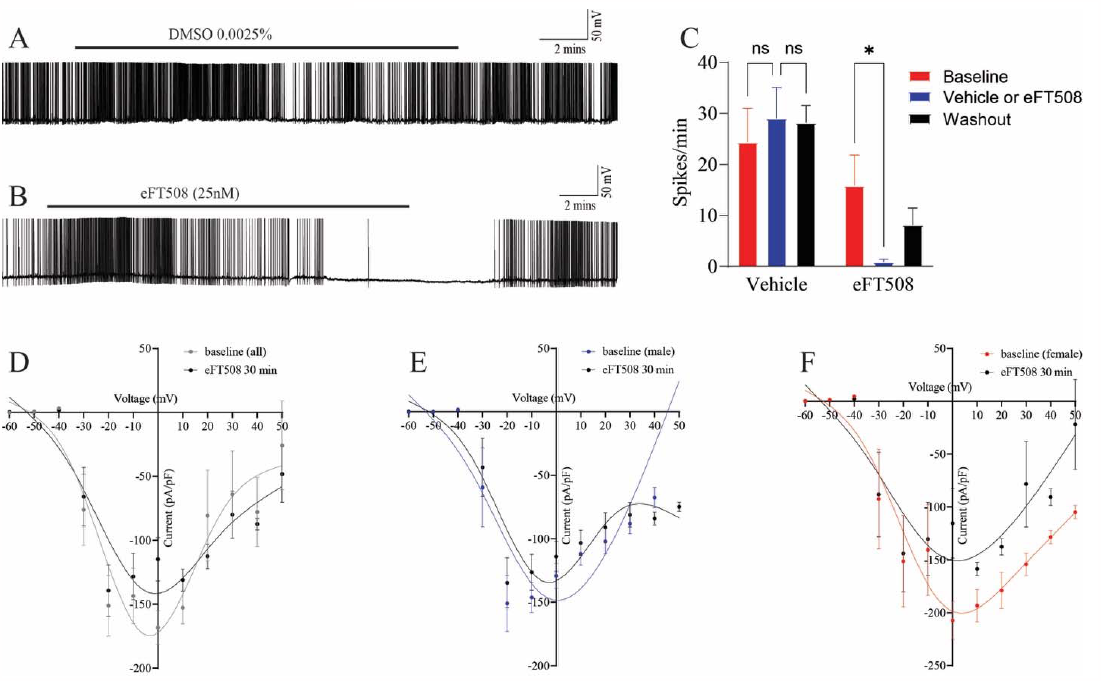
eFT508 treatment reverses SA in human DRG neurons associated with painful dermatomes. Representative recording of a human DRG neuron with SA is shown in **A** that is unaffected by vehicle treatment and a similar recording where SA is suppressed by bath application of eFT508 (25 nM) is shown in **B. C**) Combined results from 16 separate whole cell patch clamp experiments in human DRG neurons with SA treated with vehicle or eFT508 analyzed using two-way ANOVA with Tukey’s multiple comparison tests. The I–V curves shown were constructed from current traces before and after the 30-minute eFT508 (25 nM) application for all neurons (**D**), those from male patients (**E**), and those from female patients (**F**) and analyzed using two-way ANOVA with Bonferroni multiple comparisons tests. *p <0.05.

### eFT508 alters specific aspects of action potential (AP) kinetics in spontaneously active human DRG neurons

We next assessed specific influences of eFT508 treatment (25 nM, 30 min incubation) on human DRG neurons cultured from painful dermatomes of thoracic vertebrectomy patients. In neurons that did not show SA (**Fig 2A)**, eFT508 depolarized the resting membrane potential, consistent with previous findings in mice (Moy et al., 2017), and decreased membrane afterhyperpolarization amplitude but was otherwise without effect (**Fig 2C-K, Table 1**). In neurons with SA (**Fig 2B)**, eFT508 did not influence the resting membrane potential, current threshold or AP threshold but did reduce the AP amplitude, AP overshoot and reduced the magnitude of afterhyperpolarization amplitude (**Fig 2C-H, Table 1**). AP rise time and fall time were unaffected by eFT508 treatment in SA neurons although AP rise time was increased by eFT508 treatment in neurons without SA (**Fig 2I-K, Table 1**). The AP width, which is largely reflective of Nav1.8 currents in human DRG neurons (Han et al., 2015), was reduced by eFT508 treatment, although this effect was not significant due to the large variation in AP width in neurons with SA (**Fig 2K, Table 1**). The variance before and after eFT508 treatment was significantly different (p < 0.0001) by F test (F(7,7) = 72.63).

**Figure 2.**
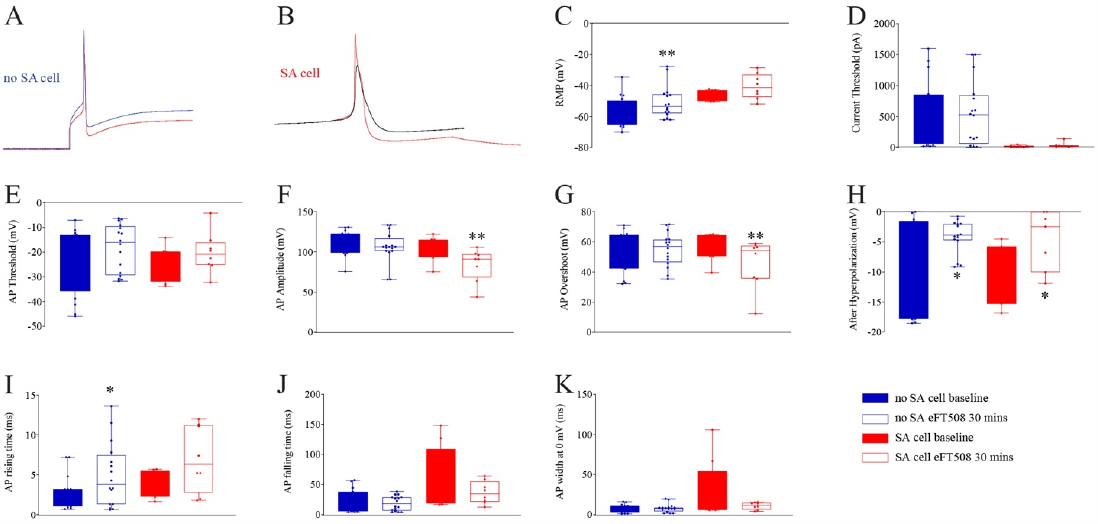
Effect of eFT508 on AP characteristics in neurons with and without SA. Neurons were divided into 2 subgroups based on whether or not the cell showed SA at baseline and compared each using non-parametric Wilcoxon matched-pairs signed rank tests. Example traces of neurons without SA (**A**, red = baseline; blue = after eFT508 treatment) and with SA (**B**, red = baseline; black = after eFT508 treatment). Treatment effect of eFT508 for no SA and SA neurons is shown for resting membrane potential (RMP) (**C**), current thresholds (**D**), AP threshold (**E**), AP amplitude (**F**), AP overshoot (**G**), AP afterhyperpolarization (**H**), AP rising time (**I**), AP falling time (**J**), and AP width at 0 mV (**K**). *p <0.05, **p <0.01.

**Table 1.** Effects of eFT508 administration on Membrane Parameters of Human Peripheral Sensory Neurons

| Neuron Type |  | RMP (mV) | Current Threshold (pA) | AP Threshold (mV) | Amplitude (mV) | Overshoot (mV) | AHP Amplitude (mV) | Rise time (ms) | Fall time (ms) | AP width at 0 mV (ms) |
| --- | --- | --- | --- | --- | --- | --- | --- | --- | --- | --- |
| No SA<br>(n = 16) | Baseline | -55.5 ± 2.4 | 543.1 ± 131.8 | -23.1 ± 3.2 | 111.0 ± 3.8 | 54.2 ± 3.8 | -7.9 ± 1.9 | 2.8 ± 0.5 | 21.1 ± 4.8 | 7.2 ± 1.3 |
|  | eFT508 30 min | -50.7 ± 2.6** | 544.4 ± 131.6 | -18.3 ± 2.4 | 106.0 ± 4.9 | 55.1 ± 2.8 | -4.1 ± 0.7* | 4.9 ± 1.0* | 18.8 ± 3.1 | 8.0 ± 1.3 |
| SA<br>(n = 8) | Baseline | -46.8 ± 1.2 | 18.8 ± 5.8 | -25.3 ± 2.4 | 103.9 ± 5.5 | 55.8 ± 3.1 | -10.4 ± 1.8 | 4.0 ± 0.6 | 55.8 ± 18.5 | 28.3 ± 13.2 |
|  | eFT508 30 min | -40.5 ± 2.9 | 36.3 ± 15.2 | -20.3 ± 2.9 | 83.9 ± 7.3** | 45.8 ± 5.8** | -4.6 ± 1.9* | 7.0 ± 1.5 | 36.7 ± 6.6 | 10.9 ± 1.6 |

Values are given as mean ± SEM. RMP: resting membrane potential; AP: action potential; Current threshold: injected current to induce action potential; AHP: after hyperpolarization amplitude
\* $p < 0.05$ , \*\* $p < 0.01$ , eFT508 vs baseline.

| Neuron Type |  | RMP (mV) | Current Thresold (pA) | AP Threshold (mV) | Amplitude (mV) | Overshoot (mV) | AHP Amplitude (mV) | Rise time (ms) | Fall time (ms) | AP width at 0 mV (ms) |
| --- | --- | --- | --- | --- | --- | --- | --- | --- | --- | --- |
| **No SA**<br>(n = 16) | Baseline | -55.5 ± 2.4 | 543.1 ± 131.8 | -23.1 ± 3.2 | 111.0 ± 3.8 | 54.2 ± 3.8 | -7.9 ± 1.9 | 2.8 ± 0.5 | 21.1 ± 4.8 | 7.2 ± 1.3 |
|  | eFT508 30 min | -50.7 ± 2.6** | 544.4 ± 131.6 | -18.3 ± 2.4 | 106.0 ± 4.9 | 55.1 ± 2.8 | -4.1 ± 0.7* | 4.9 ± 1.0* | 18.8 ± 3.1 | 8.0 ± 1.3 |
| **SA**<br>(n = 8) | Baseline | -46.8 ± 1.2 | 18.8 ± 5.8 | -25.3 ± 2.4 | 103.9 ± 5.5 | 55.8 ± 3.1 | -10.4 ± 1.8 | 4.0 ± 0.6 | 55.8 ± 18.5 | 28.3 ± 13.2 |
|  | eFT508 30 min | -40.5 ± 2.9 | 36.3 ± 15.2 | -20.3 ± 2.9 | 83.9 ± 7.3** | 45.8 ± 5.8** | -4.6 ± 1.9* | 7.0 ± 1.5 | 36.7 ± 6.6 | 10.9 ± 1.6 |

Values are given as mean ± SEM. RMP: resting membrane potential; AP: action potential; Current threshold: injected current to induce action potential; AHP: after hyperpolarization amplitude
\* $p < 0.05$ , \*\* $p < 0.01$ , eFT508 vs baseline.

### eFT508 rapidly alters eIF4E phosphorylation in human DRG neurons

MNK phosphorylates eIF4E with high specificity at S209 (Ueda et al., 2004) making this biochemical marker a precise measure of MNK activity. Effects of eFT508 on DRG neuron excitability emerge within 5 min in physiological recordings (**Fig 1**). Our biochemical assessment of MNK-mediated phosphorylation of eIF4E in human DRG neurons shows that eFT508 (25 nM) profoundly reduces eIF4E phosphorylation as soon as 2 min after treatment, peaking at 5 min after treatment with a sustained loss of 75-80% of eIF4E phosphorylation (**Fig 3A-B**).

**Figure 3.**
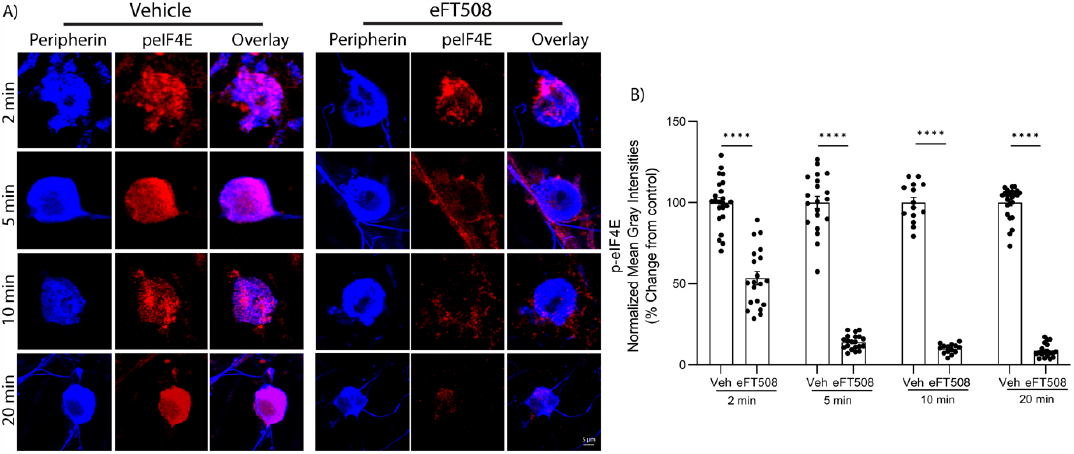
MNK inhibition with eFT508 rapidly decreases phosphorylated eIF4E in human DRG neurons. **A**) Representative zoomed-in images of p-eIF4E immunostaining (red) and neuronal marker, peripherin (Blue). Zoomed-in image scale bar: 5 μm. **B**) Normalized mean gray intensity values of p-eIF4E signal shown starting at 2 min through 20-min post eFT508 treatment. n = 23, and 20 neurons for Veh and 25nM eFT508 (2 min); n = 20, and 20 neurons for Veh and 25nM eFT508 (5 min); n = 14, and 14 neurons for Veh and 25nM eFT508 (10 min); n = 24, and 19 neurons for Veh and 25nM eFT508 (20 min). Two-way ANOVA with Bonferroni’s multiple comparisons (****P < 0.0001).

## Discussion

The major conclusion emerging from these experiments is that inhibition of MNK signaling reduces spontaneous activity in human DRG neurons recovered from thoracic vertebrectomy surgery patients. Because these neurons were recovered from DRGs innervating dermatomes associated with radicular pain, the SA observed in our experiments is almost certainly directly related to the neuropathic pain experienced by these patients prior to surgery (North et al., 2019; Ray et al., 2023). We take this as strong translational evidence that MNK inhibitors have the potential to have a profound effect on neuropathic pain in humans suffering from this disorder because they are likely to reduce the SA that is the cause of their pain.

Our work raises several important questions for future experiments. The most obvious is how does MNK inhibition reduce SA. Some important clues emerge from our recordings. The first is that while macroscopic Na^+^ and K^+^ currents were not changed in IV curves with eFT508 treatment, some specific aspects of the AP were changed with eFT508 treatment in SA neurons. One of these was AP amplitude and overshoot, both of which were reduced. Because the majority of the AP current after reaching threshold is carried by Nav1.8 in nociceptors in rats and humans, including AP overshoot (Djouhri et al., 2003; Han et al., 2015), this could be interpreted as a decrease in Nav1.8 availability following eFT508 treatment. The effects on AP width can also be interpreted as an effect on Nav1.8 currents by eFT508 treatment. A second effect was on the AP afterhyperpolarization which was decreased by eFT508 treatment in SA neurons. Afterhyperpolarization is a key driver of activation of hyperpolarization-activated cyclic nucleotide-modulated (HCN) ion channels which have been show to play a critical role in neuropathic pain in animal models (Emery et al., 2011; Tsantoulas et al., 2016). These channels are responsible for rebound depolarizations in response to hyperpolarization that is augmented by the presence of cyclic nucleotides (Tsantoulas et al., 2016), which would also be expected to be increased in nociceptors in the presence of neuropathic pain (Walters et al., 2023). Another possibility is that the decrease in afterhyperpolarizations produced by eFT508 treatment reflect decreased Ca^2+^-activated K^+^ currents that have been shown to play a critical role in rhythmic firing of neurons (Llinas, 1990; Choi et al., 2010). Decreasing the size of the afterhyperpolarization would potentially reflect decreased intracellular Ca^2+^, but it would also result in smaller recovery from inactivation for low threshold voltage-gated Ca^2+^ channels that are known to contribute to DRG neuron excitability (Waxman and Zamponi, 2014). Therefore, decreased availability of Nav1.8 and decreased recruitment of HCN and/or Ca^2+^ channels via reduced afterhyperpolarization produces a testable set of hypotheses for future experiments in these very difficult to obtain clinical samples.

Another question emerging from this work is whether these effects of MNK are mediated by translation regulation via eIF4E or via other phosphorylation targets. First, the action of eFT508 at 25 nM in the experiments we describe is very likely specific for MNK because this drug has been tested against a broad variety of off-targets and has no known other activities at the concentration we used (Stumpf et al., 2018). While the kinetics of activity-dependent translation have not been measured in human nociceptors, detailed experiments have been done in rodent brain showing that changes in the synaptic proteome can be induced through translation regulation on the order of single minutes (Hafner et al., 2019; Latallo et al., 2019). These changes have an impact on synaptic transmission suggesting that very rapid changes in protein synthesis can lead to a direct impact on neuronal excitability (Biever et al., 2019). The proteins that are synthesized are likely proteins that regulate the insertion or removal of ion channels in the plasma membrane as these proteins tend to be shorter, and do not have transmembrane domains that require compartmentalized protein synthesis machinery for proper assembly (Biever et al., 2019). Hence, one possibility is that MNK-eIF4E signaling regulates the continuous translation of mRNAs encoding proteins that regulate membrane trafficking of Nav1.8 and K^+^ channels involved in afterhyperpolarization or, potentially, HCN channels. Based on our physiology findings, we would expect all of these proteins to be removed from the plasma membrane or be placed in a state that decreases their availability to open upon MNK inhibition. Another possibility is that MNK phosphorylates ion channels or ion channel associated proteins to generate SA and that MNK inhibition rapidly reverses this effect, similar to our results for the kinase’s canonical target, eIF4E. Additional phosphorylation targets for MNK have been described, including Syngap1 which was recently discovered in a phosphoproteomic screen with MNK inhibition in brain neurons (Chalkiadaki et al., 2022). Similar phosphoproteomic screens for MNK targets in human DRG neurons could yield fruitful additional targets to better understand how MNK inhibition reduces SA in human DRG neurons.

Our work provides a clear demonstration of the utility of MNK inhibitors for reducing sensory neuron SA, providing an unprecedented level of insight into their effect on human DRG neurons associated directly with neuropathic pain in a patient population. There is clearly more to be learned about the mechanism of action of MNK inhibitors for reducing SA in human DRG neurons. However, our findings lead to testable hypotheses to better understand these mechanisms, and a strong rationale to test MNK inhibitors for neuropathic pain in clinical trials.

## Acknowledgements

The authors are grateful to the patients who participated in this study and to the organ donor and their family.

## References

Barragan-Iglesias P, Franco-Enzastiga U, Jeevakumar V, Shiers S, Wangzhou A, Granados-Soto V, Campbell ZT, Dussor G, Price TJ (2020) Type I Interferons Act Directly on Nociceptors to Produce Pain Sensitization: Implications for Viral Infection-Induced Pain. J Neurosci 40:3517–3532.

Biever A, Donlin-Asp PG, Schuman EM (2019) Local translation in neuronal processes. Curr Opin Neurobiol 57:141–148.

Chalkiadaki K, Hooshmandi M, Lach G, Statoulla E, Simbriger K, Amorim IS, Kouloulia S, Zafeiri M, Pothos P, Bonneil E, Gantois I, Popic J, Kim SH, Wong C, Cao R, Komiyama NH, Atlasi Y, Jafarnejad SM, Khoutorsky A, Gkogkas CG (2022) Mnk1/2 kinases regulate memory and autism-related behaviours via Syngap1. Brain.

Choi S, Yu E, Kim D, Urbano FJ, Makarenko V, Shin HS, Llinas RR (2010) Subthreshold membrane potential oscillations in inferior olive neurons are dynamically regulated by P/Q- and T-type calcium channels: a study in mutant mice. J Physiol 588:3031–3043.

Djouhri L, Fang X, Okuse K, Wood JN, Berry CM, Lawson SN (2003) The TTX-resistant sodium channel Nav1.8 (SNS/PN3): expression and correlation with membrane properties in rat nociceptive primary afferent neurons. J Physiol 550:739–752.

Emery EC, Young GT, Berrocoso EM, Chen L, McNaughton PA (2011) HCN2 ion channels play a central role in inflammatory and neuropathic pain. Science 333:1462–1466.

Hafner AS, Donlin-Asp PG, Leitch B, Herzog E, Schuman EM (2019) Local protein synthesis is a ubiquitous feature of neuronal pre- and postsynaptic compartments. Science 364.

Han C, Estacion M, Huang J, Vasylyev D, Zhao P, Dib-Hajj SD, Waxman SG (2015) Human Na(v)1.8: enhanced persistent and ramp currents contribute to distinct firing properties of human DRG neurons. J Neurophysiol 113:3172–3185.

Haroutounian S, Nikolajsen L, Bendtsen TF, Finnerup NB, Kristensen AD, Hasselstrom JB, Jensen TS (2014) Primary afferent input critical for maintaining spontaneous pain in peripheral neuropathy. Pain 155:1272–1279.

Lackovic J, Price TJ, Dussor G (2023) MNK1/2 contributes to periorbital hypersensitivity and hyperalgesic priming in preclinical migraine models. Brain 146:448–454.

Latallo MJ, Livingston NM, Wu B (2019) Translation imaging of single mRNAs in established cell lines and primary cultured neurons. Methods 162-163:12–22.

Li Y, North RY, Rhines LD, Tatsui CE, Rao G, Edwards DD, Cassidy RM, Harrison DS, Johansson CA, Zhang H, Dougherty PM (2018) DRG Voltage-Gated Sodium Channel 1.7 Is Upregulated in Paclitaxel-Induced Neuropathy in Rats and in Humans with Neuropathic Pain. J Neurosci 38:1124–1136.

Li Y, Adamek P, Zhang H, Tatsui CE, Rhines LD, Mrozkova P, Li Q, Kosturakis AK, Cassidy RM, Harrison DS, Cata JP, Sapire K, Zhang H, Kennamer-Chapman RM, Jawad AB, Ghetti A, Yan J, Palecek J, Dougherty PM (2015) The cancer chemotherapeutic paclitaxel increases human and rodent sensory neuron responses to TRPV1 by activation of TLR4. J Neurosci 35:13487–13500.

Llinas R (1990) Intrinsic electrical properties of nerve cells and their role in network oscillation. Cold Spring Harb Symp Quant Biol 55:933–938.

Megat S, Ray PR, Moy JK, Lou TF, Barragan-Iglesias P, Li Y, Pradhan G, Wanghzou A, Ahmad A, Burton MD, North RY, Dougherty PM, Khoutorsky A, Sonenberg N, Webster KR, Dussor G, Campbell ZT, Price TJ (2019) Nociceptor Translational Profiling Reveals the Ragulator-Rag GTPase Complex as a Critical Generator of Neuropathic Pain. J Neurosci 39:393–411.

Mihail SM, Wangzhou A, Kunjilwar KK, Moy JK, Dussor G, Walters ET, Price TJ (2019) MNK-eIF4E signalling is a highly conserved mechanism for sensory neuron axonal plasticity: evidence from Aplysia californica. Philos Trans R Soc Lond B Biol Sci 374:20190289.

Moy JK, Khoutorsky A, Asiedu MN, Black BJ, Kuhn JL, Barragan-Iglesias P, Megat S, Burton MD, Burgos-Vega CC, Melemedjian OK, Boitano S, Vagner J, Gkogkas CG, Pancrazio JJ, Mogil JS, Dussor G, Sonenberg N, Price TJ (2017) The MNK-eIF4E Signaling Axis Contributes to Injury-Induced Nociceptive Plasticity and the Development of Chronic Pain. J Neurosci 37:7481–7499.

North RY, Li Y, Ray P, Rhines LD, Tatsui CE, Rao G, Johansson CA, Zhang H, Kim YH, Zhang B, Dussor G, Kim TH, Price TJ, Dougherty PM (2019) Electrophysiological and transcriptomic correlates of neuropathic pain in human dorsal root ganglion neurons. Brain 142:1215–1226.

Ray PR, Shiers S, Caruso JP, Tavares-Ferreira D, Sankaranarayanan I, Uhelski ML, Li Y, North RY, Tatsui C, Dussor G, Burton MD, Dougherty PM, Price TJ (2023) RNA profiling of human dorsal root ganglia reveals sex differences in mechanisms promoting neuropathic pain. Brain 146:749–766.

Serra J, Bostock H, Sola R, Aleu J, Garcia E, Cokic B, Navarro X, Quiles C (2012) Microneurographic identification of spontaneous activity in C-nociceptors in neuropathic pain states in humans and rats. Pain 153:42–55.

Shiers S, Mwirigi J, Pradhan G, Kume M, Black B, Barragan-Iglesias P, Moy JK, Dussor G, Pancrazio JJ, Kroener S, Price TJ (2020) Reversal of peripheral nerve injury-induced neuropathic pain and cognitive dysfunction via genetic and tomivosertib targeting of MNK. Neuropsychopharmacology 45:524–533.

Stumpf CR, Chen J, Goel VK, Parker GS, Thompson PA, Webster KR (2018) Inhibition of MNK by eFT508 reprograms T-cell signaling to promote an anti-tumor immune response. In: American Association for Cancer Research. Chicago, IL.

Tsantoulas C, Mooney ER, McNaughton PA (2016) HCN2 ion channels: basic science opens up possibilities for therapeutic intervention in neuropathic pain. Biochem J 473:2717–2736.

Ueda T, Watanabe-Fukunaga R, Fukuyama H, Nagata S, Fukunaga R (2004) Mnk2 and Mnk1 are essential for constitutive and inducible phosphorylation of eukaryotic initiation factor 4E but not for cell growth or development. Mol Cell Biol 24:6539–6549.

Valtcheva MV, Copits BA, Davidson S, Sheahan TD, Pullen MY, McCall JG, Dikranian K, Gereau IV RW (2016) Surgical extraction of human dorsal root ganglia from organ donors and preparation of primary sensory neuron cultures. Nature protocols 11:1877–1888.

Walters ET, Crook RJ, Neely GG, Price TJ, Smith ESJ (2023) Persistent nociceptor hyperactivity as a painful evolutionary adaptation. Trends Neurosci 46:211–227.

Waxman SG, Zamponi GW (2014) Regulating excitability of peripheral afferents: emerging ion channel targets. Nat Neurosci 17:153–163.

